# Maf1 Cooperates with Progesterone Receptor to Repress RNA Polymerase III Transcription of Select tRNAs

**DOI:** 10.1101/2024.12.16.628719

**Authors:** Jessica Finlay-Schultz, Kiran V. Paul, Benjamin Erickson, Lynsey M. Fettig, Benjamin S. Hastings, Deborah L. Johnson, David L. Bentley, Peter Kabos, Carol A. Sartorius

## Abstract

Progesterone receptors (PR) can regulate transcription by RNA Polymerase III (Pol III), which transcribes small non-coding RNAs, including all transfer RNAs (tRNAs). We have previously demonstrated that PR is associated with the Pol III complex at tRNA genes and that progestins downregulate tRNA transcripts in breast tumor models. To further elucidate the mechanism of PR-mediated regulation of Pol III, we studied the interplay between PR, the Pol III repressor Maf1, and TFIIIB, a core transcription component. ChIP-seq was performed for PR, the Pol III subunit POLR3A, the TFIIIB component Brf1, and Maf1 in breast cancer cells with or without progestin treatment. Upon progestin exposure, PR localized to approximately half of POLR3A-occupied tRNA genes, with Maf1 co-recruited to many of these PR-POLR3A sites. While progestin treatment did not significantly alter the number of tRNA genes occupied by Pol III or Brf1, Brf1 occupancy was stabilized, as indicated by increased peak amplitudes. Analysis of nascent tRNA transcription revealed a specific progestin-induced downregulation of approximately one-third of highly expressed tRNA genes. This repression was significantly reduced by Maf1 knockdown, indicating that Maf1 is necessary for PR-mediated tRNA transcription downregulation. Overall, these findings demonstrate a ligand-dependent PR-mediated repression of tRNA transcription through Maf1.

## INTRODUCTION

Progesterone receptors (PR) are crucial ligand-activated transcription factors that regulate the delicate balance between growth and differentiation in target tissues. The steroid hormone progesterone, the primary natural ligand for PR, plays a pivotal role in regulating reproductive tissue function, including the cyclical expansion of mammary stem cells within the breast. In these processes, progesterone acts in concert with 17β-estradiol, the main ligand for estrogen receptors (ER) (1–3), to coordinate hormonal signaling in reproductive organs. PR are expressed in approximately half of all breast cancers in parallel with ER and serve as biomarkers for sensitivity to endocrine therapies (4,5). Progesterone or synthetic PR-binding ligands (progestins) generally counteract the mitogenic effects of estrogens in breast cancer, although PR can alternatively cooperate with ER under certain conditions (6,7). This crosstalk is believed to modulate ER’s transcriptional activity, as PR and ER co-localize at numerous genomic sites under estrogenic conditions, with progestins redirecting ER and PR to an alternative cistrome (8,9). In our prior investigations using PR+ breast tumor models subjected to prolonged progestin treatment, we identified a significant association between PR and RNA polymerase III (Pol III) regulated tRNA genes, resulting in the downregulation of both pre- and mature tRNAs (10). Notably, PR regulation of Pol III activity appears to not involve direct ER association at these loci, suggesting a mechanism whereby PR influences Pol III-mediated transcription independently of direct ER interactions.

Pol III is responsible for transcribing essential small non-coding RNAs crucial for translation, such as 5S rRNA and all tRNAs, and its dysregulation is commonly observed in cancer (11). Oncogenes like c-Myc and PI3K/mTOR typically enhance Pol III activity, while tumor suppressors such as p53, RB, and PTEN reduce Pol III activity (12,13). Additionally, Pol III activity is tightly regulated by alterations in the core transcriptional complexes TFIIIB and TFIIIC, along with the conserved repressor Maf1. The rate limiting TFIIIB complex, comprising Brf1, TATA-binding protein (TBP), and Bdp1, is often upregulated in various cancers, including ER+ breast, ovarian, liver, and lung cancers (14–18). Maf1, a key Pol III repressor, is evolutionarily conserved from yeast to humans, and acts as a tumor suppressor (19–22). In human cells, Maf1 is predominantly found in both the cytoplasm and nucleus under basal conditions and is phosphorylated by mTOR signaling, which inactivates Maf1; inhibition of mTOR results in Maf1 dephosphorylation and subsequent Pol III inhibition (20,21,23). Studies on regulation of Maf1 and its activity in breast cancer are limited, but recent research suggests its association with positive responses to HER2-targeted therapies (24). Considering breast cancer is strongly influenced by steroid hormones, it serves as an ideal model system to understand how progesterone signaling affects Pol III transcription, revealing novel aspects of PR’s contribution to breast cancer cell growth.

Here we demonstrate that short-term progestin treatment induces co-occupancy of PR and Maf1 at a subset of tRNA genes in PR-positive breast cancer cells. PR-targeted tRNA genes exhibit a rapid reduction in pre-tRNA transcription, a process impaired by Maf1 depletion. Notably, Brf1 localization relative to POLR3A at tRNA genes increased with progestin treatment. This study supports the idea that activated PR modulates Pol III activity by engaging Maf1 and repressing transcription without displacing Pol III from the complex. As alterations in tRNA gene expression are key determinants for cell growth and an oncogenic state, these results provide insights into the role of progestins in breast cancer growth through its ability to modulate tRNA gene activity and uncovers a novel mechanism for Pol III-mediated transcription regulation.

## MATERIALS AND METHODS

### Reagents

Hormones (progesterone (P4), promegestone (R5020), and 17β-estradiol (E2)) were purchased from MilliporeSigma (St. Louis, MO, USA).

### Biological Resources

T47D breast cancer cells were obtained from the University of Colorado Cancer Center Cell Technologies Shared Resource. UCD4 breast cancer cells were developed by our laboratory as described (25). T47D cells were maintained in minimal Eagle’s medium, 5% fetal bovine serum (FBS), 1X NEAA, 1×10^-9^ M insulin, 0.1 mg/mL sodium pyruvate, and 2 mM L-glutamine. UCD4 cells were maintained in DMEM/F12 supplemented with 10% FBS, 100 ng/mL choleratoxin, and 10^−9^ M insulin. For all experiments using the UCD4 line, cells were pretreated with 10 nM E2 for 48 h to induce PR expression. Cell lines were authenticated using short tandem repeat (STR) analysis and tested negative for mycoplasma using the MycoAlert mycoplasma detection kit (Lonza, Basel, Switzerland). Sigma Mission shRNAs targeting Maf1 (TRCN0000182935, TRCN0000180558) and a non-targeting clone (SHC0002) were purchased from the University of Colorado Functional Genomics Shared Resource. Cells were transduced with virus containing the shRNAs and stable pools selected with puromycin.

### qPCR

Total RNA was extracted using QIAzol lysis reagent and the miRNeasy kit (Qiagen, Venlo, Netherlands). cDNA was prepared using the Verso cDNA synthesis kit (ThermoFisher). qRT-PCR was performed on cDNA or ChIP DNA using Absolute Blue Sybr Green Low ROX (ThermoFisher). Analysis was performed using the Pfaffl method (26). All primers used for these studies are found in Supplementary Table 1.

### ChIP-seq

T47D cells were grown to 70-80% confluency in 15 cm dishes in phenol red free media containing charcoal stripped serum and treated with 100 nM P4, 10 nM R5020, or vehicle for 1 hour. Immunoprecipitation was performed using antibody PR Ab-8 (MS-298-P, ThermoFisher, Grand Island, NY, USA), PR F-4 (sc-7208, Santa Cruz Biotechnology, Dalla, TX, USA), POLR3A D5Y2D (12825, Cell Signaling Technology, Danvers, MA, USA), Brf1 (A301-228A, Bethyl Laboratories, Montgomery, TX, USA), Maf1 (GTX107667, GeneTex, Irvine, CA, USA), or a corresponding IgG negative control of the same species. For data in Figure 1, cells were processed using the ChIP-IT Express kit (Active Motif, Carlsbad, CA, USA) with chromatin sheared using a S220 Focused Ultrasonicator (Covaris, Woburn, MA, USA). For data in Figure 2, cells were processed as described (27) using methanol-free formaldehyde with chromatin sheared using an ME220 Focused Ultrasonicator (Covaris). DNA was purified and used for qPCR or library preparation. Library preparation for sequencing experiments was performed for Figure 1 using the Illumina TruSeq ChIP-Seq Library Preparation kit (Illumina, San Diego, CA, USA) and for Figure 2 using the NEBNext Ultra II DNA Library Prep Kit for Illumina (New England Biolabs, Ipswich, MA, USA). Input DNA was also used as a control. ChIP-seq paired-end reads were processed with cutadapt (https://cutadapt.readthedocs.io/en/stable/index.html) to remove 3’ adaptor sequences and 3’ bases with QUAL < 10. Trimmed reads were aligned to the human genome (GRCh38/hg38 build) using bowtie2 (https://bowtie-bio.sourceforge.net/bowtie2/index.shtml). Peaks were called on aligned reads using MACS2 callpeak (https://hbctraining.github.io/Intro-to-ChIPseq/lessons/05_peak_calling_macs.html) which is a model-based analysis tool to determine the significance of enriched ChIP regions. Bedtools intersect was used to search for tRNA and mRNA sites in the peak regions (https://bedtools.readthedocs.io/en/latest/content/tools/intersect.html).The called peaks with tRNA and mRNA sites were then visualized using deeptools (https://deeptools.readthedocs.io/en/latest/) and custom-made scripts. Motif discovery and enrichment analysis were performed using MEME suite (https://meme-suite.org/meme/). Raw sequence data and peak calls have been deposited in the Gene Expression Omnibus (GEO) database (GSE283978, GSE283979).

**Figure 1.**
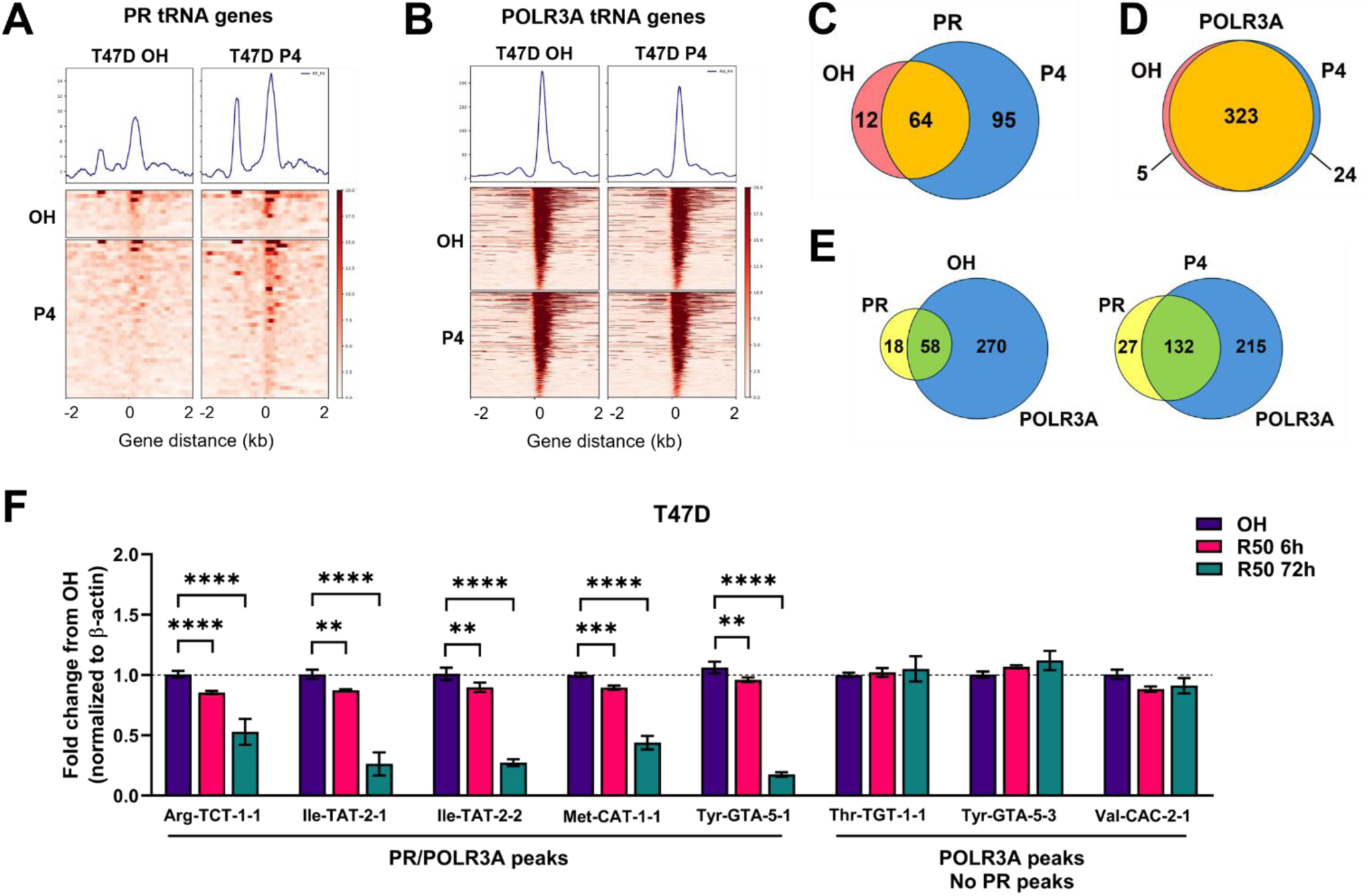
Progestin treatment increases PR recruitment to POLR3A-occupied tRNA genes and downregulates pre-tRNAs. Average signal plots and heatmaps for **(A)** PR ChIP-seq and **(B)** POLR3A ChIP-seq binding events at tRNA genes in a horizontal window of +/− 2 kb in T47D cells treated with ethanol vehicle (OH) or 100 nM P4 for 1 h. Venn diagrams depict overlap between ChIP-seq binding events at tRNA genes for **(C)** PR with OH vs P4 treatment, **(D)** POLR3A with OH vs P4 treatment, and **(E)** PR and POLR3A with OH vs P4 treatment. **(F)** qPCR analysis of pre-tRNA transcripts in T47D cells treated with vehicle (OH) or 10 nM R50 for 6 or 72 h. At least six biological replicates were used for analysis. Error bars depict mean +/−SEM. Significance was determined using Student’s t test. * p<0.05, ** p<0.01, *** p<0.001, **** p<0.0001.

**Figure 2.**
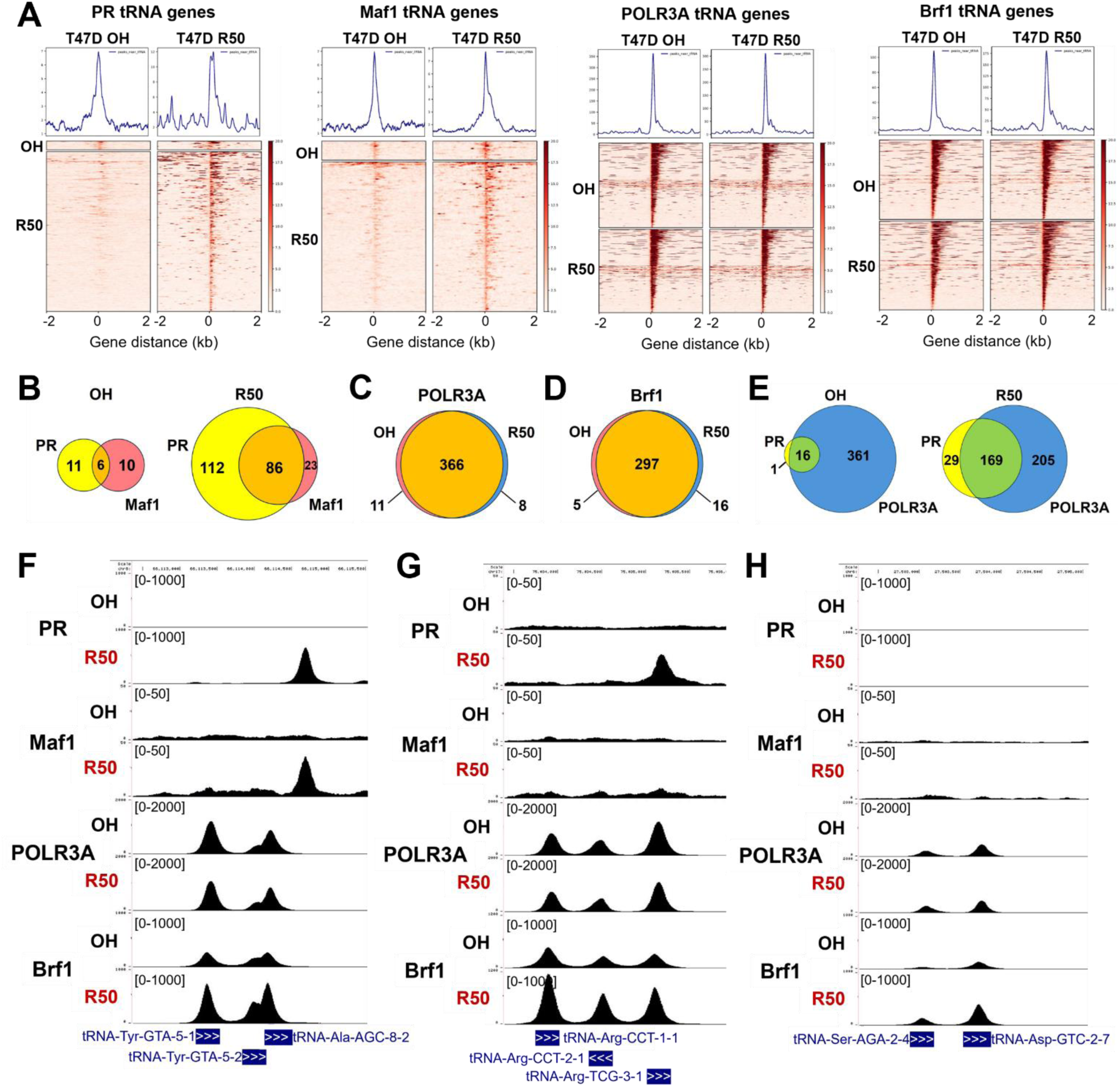
Maf1 is recruited to select PR occupied tRNA genes with progestin treatment. **(A)** Average signal plots and heatmaps for ChIP-seq tRNA binding events with antibodies to PR, Maf1, POLR3A, and Brf1 in T47D cells treated with ethanol vehicle (OH) or 10 nM R5020 (R50) for 1 h. Experiments were in biological triplicates. Heatmaps are in a horizontal window of +/− 2 kb. Venn diagrams depict overlap between ChIP-seq binding events at 429 tRNA genes from tRNAscan-SE with OH and R5020 (R50) treatment for **(B)** PR and Maf1, **(C)** POLR3A, **(D)** Brf1, and **(E)** PR and POLR3A. Representative peak tracks for each factor at tRNA genes with ethanol (OH) or R5020 (R50) depicting **(F)** PR and Maf1 recruitment, **(G)** PR but not Maf1 recruitment, and (**H**) no PR or Maf1 recruitment.

### Bromouridine sequencing (Bru-Seq)

Nascent RNA analysis by Bru-Seq was as described (28) with minor modifications. T47D cells were treated with OH or 10 nM R5020 for 1h, then labelled for 30 min with 2 mM Bromouridine (850187, MilliporeSigma). RNA (50 mg) was fragmented by ZnCl2 treatment (10mM, 70C 8 min, stopped with 10 mM EDTA) and immunoprecipitated in IP buffer (1x PBS, 0.05% Triton X-100, 1 mM DTT, RNase inhibitor) for 1 hr at 4C with 3D4 monoclonal anti-BrdU (3 μg), immobilized on protein G Dynabeads (ThermoFisher). Beads were washed 3X in IP buffer and RNA was eluted at 95C for 3 min in 5mM DTT. RNA-seq libraries were made with the KAPA RNA HyperPrep kit (Roche Diagnostics Corporation, Indianapolis, IN, USA). Reads were mapped to the hg38 UCSC human genome with Bowtie2 version 2.3.2. PCR duplicates were removed using bbtools clumpify and adapters were trimmed using bbtools bbkuk version 39.01. After filtering out rRNA, reads were mapped to the hg38 UCSC human genome with Bowtie2 version 2.3.2. Read counts were normalized to total counts on the mitochondrial chromosome.

### Immunocytochemistry

Immunocytochemistry was performed as described previously (29) in cells treated with ethanol (OH) vehicle or 10 nM R5020 for 1 h, using antibodies to Maf1 (Invitrogen PA5-21791) and PR (DAKO 1294) with DAPI nuclear stain. Images were merged using Adobe Photoshop 2024 with linear adjustments to brightness.

### Immunoblotting

Immunoblots were performed essentially as described (10). Whole-cell lysates were harvested in lysis buffer (50 mmol/L Tris pH 7.4, 140 nmol/L NaCl, 2 mmol/L EGTA, 1.0% Tween-20, 1x Halt protease and phosphatase inhibitor cocktail (ThermoFisher)). For cytosol and nuclear extracts, cells were treated with vehicle or 10 nM R5020 for 1 h, collected in cytosolic lysis buffer pH 7.9 (10 mM HEPES, 1.5 mL MgCl2, 10 mM KCl, 0.05% NP-40, 0.5 mM DTT, 1x Halt protease and phosphatase inhibitor cocktail), incubated on ice 15 minutes, centrifuged to separate the cytosolic fraction from nuclei. The resulting nuclei were resuspended in nuclear extraction buffer pH 7.9 (5 mM HEPES, 300 mM NaCl, 1.5 mM MgCl2, 0.2 mM EDTA, 26% glycerol, 0.5 mM DTT, 1x Halt protease and phosphatase inhibitor cocktail), homogenized with 20 strokes in a Dounce homogenizer, incubated on ice 30 minutes, then centrifuged to remove nuclear membranes and supernatant collected for nuclear fraction. Antibodies for immunoblots were PR (Cell Signaling Technology 8757; DAKO 1294), POLR3A (Cell Signaling Technology 12825; Bio-Rad Laboratories MCA6411GA), Maf1 (Invitrogen PA5-21791; Santa Cruz sc-365312), Brf1 (Bethyl A301-228A), and alpha-tubulin (Sigma, ST1568). For imaging, the Odyssey Infrared Imaging System (Li-Cor Biosciences) was used, with secondary antibodies IRDye800CW Goat-Anti-Mouse-IgG and IRDye680LT Goat-Anti-Rabbit-IgG (Li-Cor Biosciences).

### Statistical Analyses

Statistics were performed using Graphpad Prism 10.4. Two-tailed Student’s t-tests, or one-way ANOVA followed by Tukey post hoc multiple comparison tests were used as indicated. *P*<0.05 were considered significant.

## RESULTS

### Progesterone rapidly recruits PR to POLR3A occupied tRNA genes

We previously established that PR associated at a large fraction of tRNA genes in ER and PR positive breast cancers grown under chronic estrogen (E2) or E2 plus progesterone (P4) supplementation (10). Tumors treated with E2+P4 compared to E2 alone showed significant decreases in pre-tRNA transcripts of PR-occupied tRNA genes (10). Here we tested how acute P4 treatment affected PR and Pol III localization at tRNA genes by performing ChIP-seq for PR and the Pol III subunit POLR3A in T47D breast cancer cells treated with ethanol vehicle (OH) or progesterone (P4) for 1 h. PR binding events were enriched with P4 while POLR3A binding events remained steady across the 429 annotated high-confidence tRNA genes in the human genome (Figure 1A & B) (30). The number of PR binding events near tRNA genes doubled with P4 treatment (76 to 159) (Figure 1C). POLR3A occupied >80% of tRNA genes under vehicle and P4 conditions (Figure 1D). Overlap between PR and POLR3A binding events indicates PR is recruited to a subset of POLR3A-occupied tRNAs with P4 treatment (Figure 1E). Thus, P4 treatment induces rapid PR occupancy at a subset of tRNA genes in breast cancer cells.

### Progestin treatment rapidly downregulates pre-tRNA expression

To assess the temporal impact of progestin treatment on pre-tRNA expression, we measured pre-tRNA transcript levels over several timepoints in breast cancer cells treated with the synthetic P4 analog promegestone (R5020). T47D and UCD4 cells were treated with vehicle or R5020 for 6 or 72 h and transcripts measured by qPCR (Figure 1F). We measured pre-tRNA transcripts that showed either PR and POLR3A occupancy or only POLR3A occupancy in our T47D ChIP-seq dataset. Pre-tRNA transcript levels for tRNA genes co-occupied by PR and/or POLR3A showed a modest but significant decrease after 6 h of R5020 treatment with larger reductions observed by 72 h. tRNA genes that were occupied by POLR3A but not PR showed no changes in tRNA transcripts with R5020 treatment. This was recapitulated in UCD4 cells (Supplementary Figure S1). These data support that PR occupancy at select tRNA genes correlates with rapid decreases in pre-tRNA transcription.

### PR and Maf1 co-occupy select tRNA genes in response to progestin treatment

To investigate how the Pol III transcription complex is impacted by progestins, we performed ChIP-seq for PR, POLR3A, the TFIIIB subunit Brf1, and the Pol III repressor Maf1 in T47D cells treated with vehicle or R5020 for 1 h. Consistent with previous findings from our group and others, progestin treatment significantly increased the number of genomic PR binding sites (Supplementary Figure S2A). DNA binding events at the 429 tRNA genes were evaluated for each of the four factors. Heatmaps illustrate that with R5020 treatment, PR and Maf1 binding is enriched at tRNA genes, whereas R5020 does not affect the POLR3A and Brf1 binding at tRNA loci (Figure 2A). Upon progestin treatment, PR binding was observed at nearly half of the tRNA genes (198 total) (Figure 2B), consistent with our data in Figure 1. Notably, Maf1 was co-recruited to a substantial subset of these same tRNA genes (86 genes) under progestin treatment. In contrast, the overall occupancy of POLR3A and Brf1 at tRNA genes remained largely unchanged by progestin treatment, with these factors occupying approximately 90% and 75% of tRNA loci, respectively (Figure 2C, D). As illustrated in Figure 2E, PR was recruited predominantly to POLR3A occupied tRNA genes with progestin treatment. Transcription factor occupation at tRNA genes is illustrated with peak visualization using the NCBI Genome Browser at representative tRNA genes in Figure 2F-H. The strongest associations are observed at tRNA genes where both PR and Maf1 are recruited following progestin treatment, suggesting selective co-regulation at these loci. In contrast some tRNA genes exhibit PR binding without Maf1 or with minimal PR or Maf1. Of note, genes that showed PR but not Maf1 recruitment generally had very small PR peaks, indicating that PR recruitment at these genes is very different in magnitude (compare scales of PR in Figure 2F and 2G). Progestin-activated PR was associated with tRNA genes corresponding to all 49 codons, indicating no specific preference for particular tRNA families. A summary of PR, Maf1, Brf1, and POLR3A binding at all 429 tRNA genes, organized by codon family, is provided in Supplementary Table S2.

Motif analysis was conducted to examine sequence identities near PR binding sites. Progestin treatment induced PR binding at mRNA genes near sequences resembling progesterone response elements (PREs) (Supplementary Figure S2B). However, at tRNA genes, the only significant motif detected was the tRNA promoter B-Box (Supplementary Figure S2C), suggesting a non-traditional mode of association.

### Progestin treatment enhances Brf1 binding on tRNA genes

Maf1 has been reported to repress Pol III transcription through interaction with Brf1 to block Pol III assembly at tRNA genes, and/or by directly interacting with Pol III to dissociate it from DNA (22,31,32). Our data in breast cancer cells indicate no significant changes in the number of tRNA loci occupied by Pol III and Brf1 with progestin treatment. Profile plots of our ChIP-seq data reiterate that PR and Maf1 show marked progestin-induced enrichment near tRNA genes, while POLR3A is not significantly affected while Brf1 occupancy increased with R5020 treatment (Figure 3A). To further investigate this data, we analyzed peak amplitudes at tRNA genes under vehicle and progestin conditions and plotted the amplitudes at each tRNA locus (Figure 3B). R5020 significantly increased tRNA peak amplitudes for PR and Maf1, while POLR3A showed no significant changes, and Brf1 occupancy at tRNA genes increased significantly. Notably, progestin treatment has no effect on Brf1 or Maf1 even after 24 h (Supplementary Figure S3). We then compared tRNA peak amplitudes for POLR3A (Figure 3C) and Brf1 (Figure 3D) at genes with and without co-occupancy of PR or Maf1, as identified by ChIP-seq. Interestingly, both POLR3A and Brf1 peak amplitudes were significantly higher when PR or Maf1 was present. Together, these results suggest the formation of an inhibitory complex at PR/Maf1-occupied tRNA genes, associated with stable Brf1 and POLR3A binding. This supports a novel mechanism by which Pol III-dependent transcription can be regulated.

**Figure 3.**
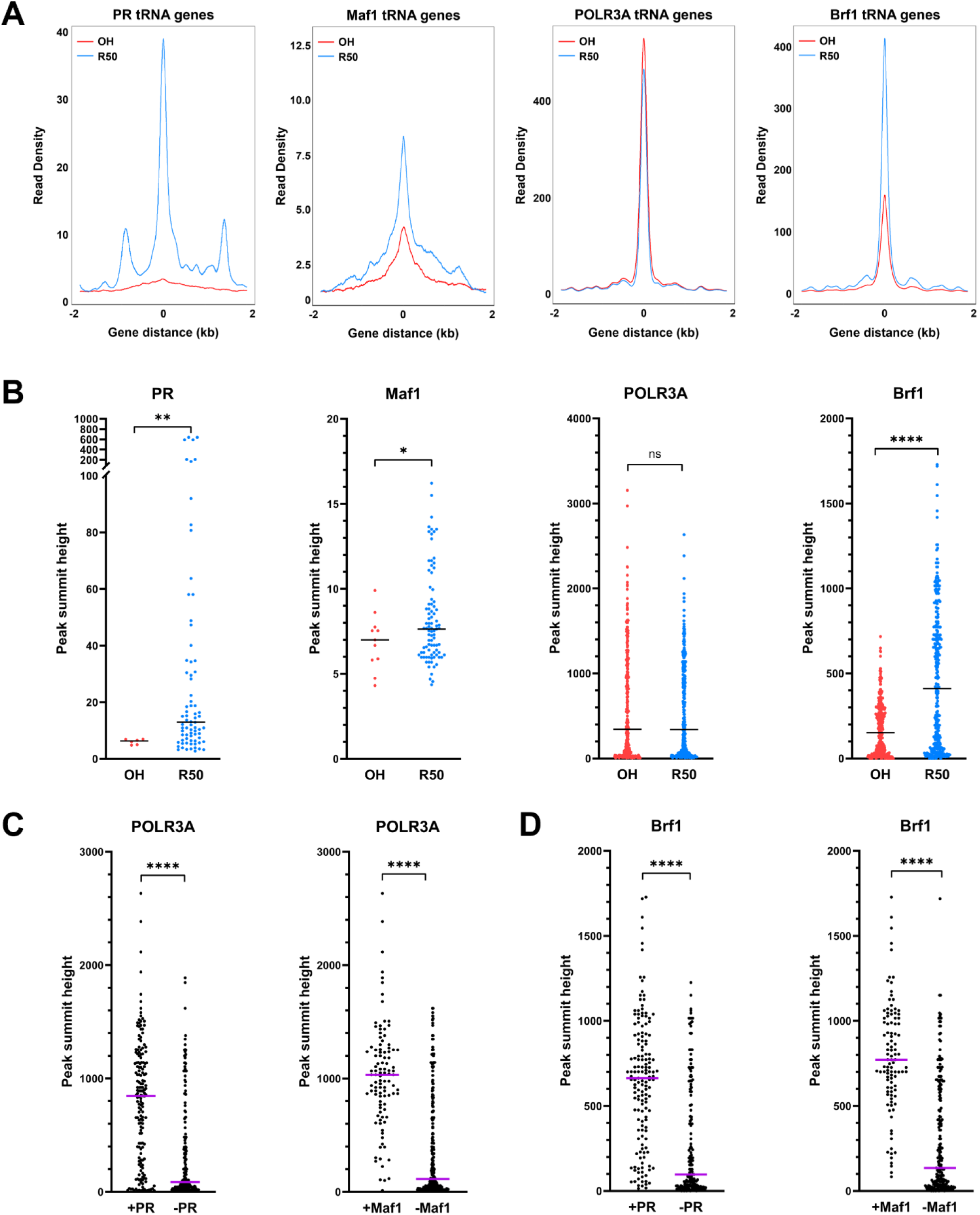
Recruitment of PR at tRNA genes stabilizes Brf1 occupancy. **(A)** Peak amplitudes (summit height) for PR, Maf1, POLR3A, and Brf1 at tRNA genes with OH or R5020 (R50) from ChIP-seq data. Peak amplitudes for **(B)** PR, Maf1, POLR3A, and Brf1 with OH or R50 graphed by individual tRNA genes. **(C)** POLR3A or **(D)** Brf1 peak amplitudes at tRNA genes with R50 with or without corresponding PR occupancy (left panels) or Maf1 (right panels). Median amplitudes are indicated. Significance was determined using Student’s t test with Welch’s correction. * p<0.05, ** p<0.01, *** p<0.001, **** p<0.0001.

### Maf1 localization is not impacted by progestin treatment

To determine whether progestins impact Maf1 cellular localization we performed ICC for PR and Maf1 without or with 1 h of R5020 treatment in T47D and UCD4 cells (Supplementary Figure S4A). PR was predominantly nuclear regardless of progestin treatment, while Maf1 remained consistently distributed between the cytoplasm and nucleus. Immunoblotting of cytosolic and nuclear fractions from T47D or UCD4 cells treated with vehicle or R5020 confirmed that PR, POLR3A, and Brf1 were predominantly localized in the nuclear fraction with or without progestin (Supplementary Figure S4B). Maf1 was present in the cytosol and nucleus, showing no significant change with progestin treatment. This indicates that R50 treatment had little effect on Maf1 localization, suggesting that progestin-induced changes in Maf1 binding occur independently of Maf1 subcellular distribution.

### Maf1 is a key factor in PR-mediated downregulation of tRNA genes

To assess how progestins influence global pre-tRNA transcription, we used Bromouridine-sequencing (Bru-seq) (28) to analyze tRNA gene transcription in T47D cells treated with either vehicle or R5020 for 1 h (Figure 4). Progestin treatment led to a reduction in mean global tRNA transcription across genes that showed POLR3A and PR binding in our ChIP-seq data (Figure 4A). This reduction is muted at genes that showed no statistical PR binding. tRNA genes with no POLR3A binding exhibited no Bru label incorporation. Pol III transcription of U6 snRNA and 7SK were unaffected by progestin treatment (Supplementary Figure S5), indicating that repression may be specific to tRNA genes. Representative Bru-seq peaks are shown in Figure 4B-D; these are matched to the ChIP-seq peaks in Figure 2F-H for ease of comparison. POLR3A-bound tRNA genes with PR and Maf1 recruitment showed R50-mediated downregulation of Bru incorporation (Figure 4B) while tRNA genes with no PR or Maf1 showed no downregulation (Figure 4D). Interestingly, the representative tRNA genes that showed PR but not Maf1 were very poorly expressed (Figure 4C); this suggests that PR may specifically target more highly-expressed tRNA genes.

**Figure 4.**
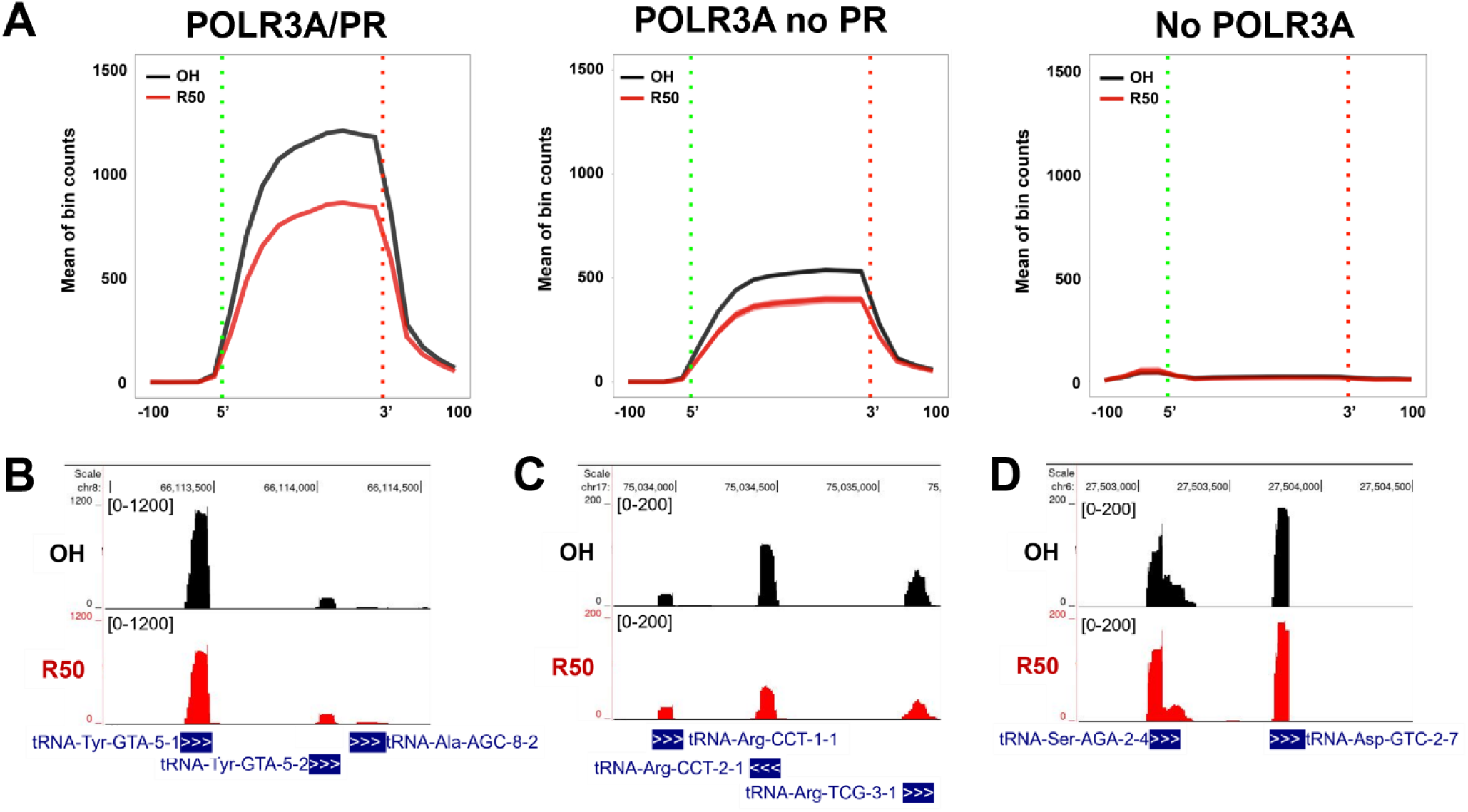
Nascent tRNA transcription rapidly decreases with progestin treatment. Bru-seq analysis in T47D cells treated with vehicle (OH) or 10 nM R5020 (R50) for 1 h followed by 30 min Bru pulse. (**A**) Relative Bru incorporation by tRNA genes occupied by POLR3A/PR, POLR3A alone, or none of the three factors from R50 ChIP-seq data. Representative peak tracks for Bru incorporation by tRNA genes with ethanol (OH) or R5020 (R50) at loci with **(B)** PR and Maf1 recruitment, **(C)** PR but not Maf1 recruitment, and (**D**) no PR or Maf1 recruitment.

To investigate the contribution of Maf1 in PR-mediated downregulation of tRNA genes, we developed T47D cells with stable Maf1 short hairpin (sh) RNA knockdown (Figure 5A). Bru-seq analysis of T47D-shControl (shCont) and shMaf1 cells following progestin treatment showed that shCont cells exhibited transcription patterns consistent with our original T47D Bru-seq data. Pie charts denote that progestins decreased transcription of over one third of tRNA genes (Figure 5B, green areas). Notably, Maf1 knockdown significantly reduced the number of progestin-downregulated tRNAs by half. The decrease in tRNA transcription was less pronounced in shMaf1 cells compared to wild type cells, both for genes with POLR3A and PR binding as well as POLR3A without PR binding (Figure 5C). Maf1 knockdown led to increased Bru incorporation at POLR3A-bound tRNA genes, indicating an overall upregulation of tRNA gene transcription (compare Figure 4A and Figure 5C). tRNA genes with no POLR3A binding exhibited no Bru label incorporation. Progestin treatment led to a reduction in mean global tRNA transcription at all high-confidence tRNA genes, with knockdown muting the reduction (Figure 5D). Overall these results suggest that Maf1 is necessary for progestin-induced tRNA downregulation.

**Figure 5.**
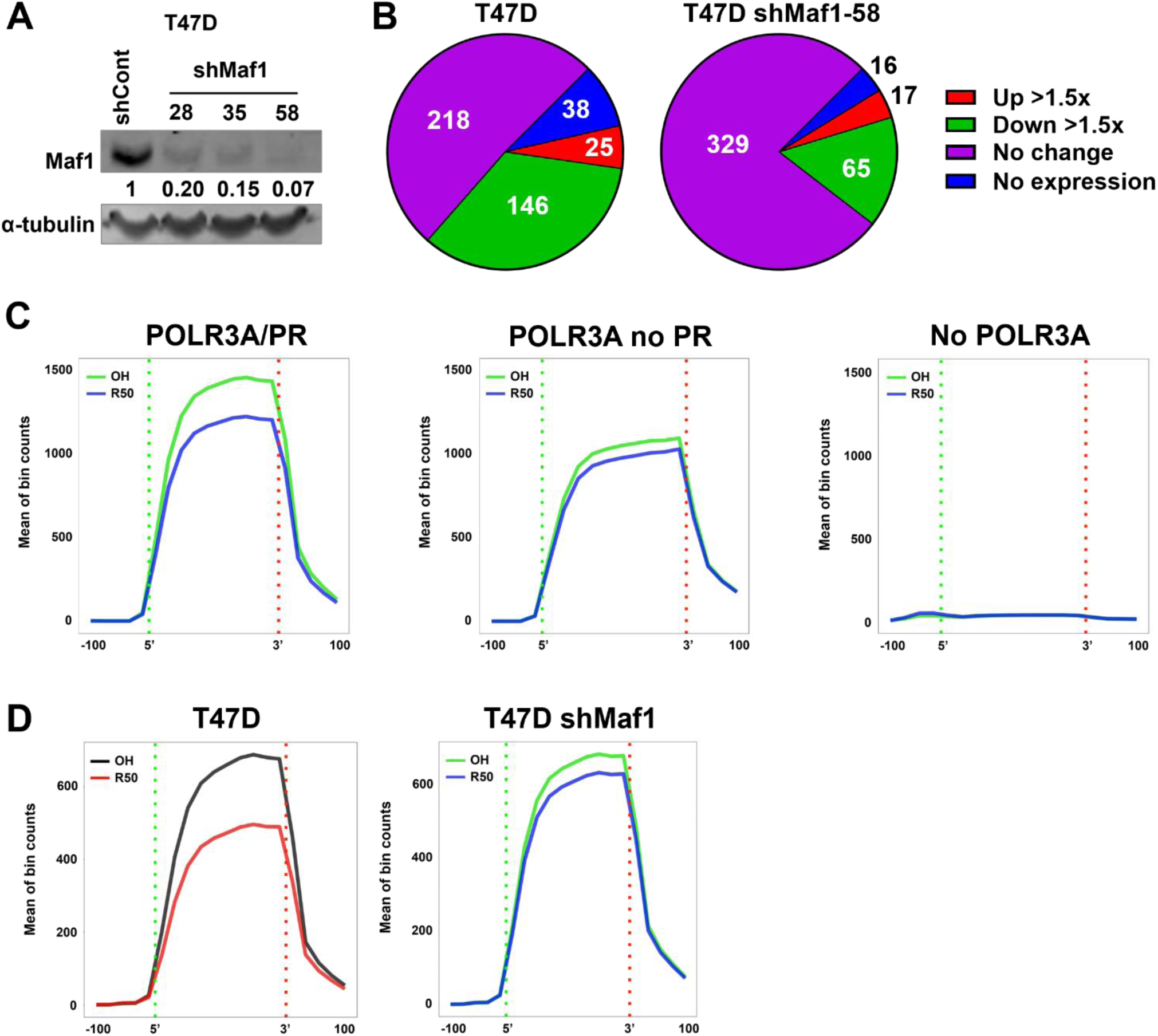
Progestin-induced nascent tRNA transcription is reduced by shMaf1. Bru-seq analysis in T47D shMaf1 cells treated with vehicle (OH) or 10 nM R5020 (R50) for 1 h followed by 30 min Bru pulse. (**A**) Immunoblot of T47D cells stably expressing shControl (shCont) and shMaf1 constructs 28, 35, and 58. Ratio of Maf1 expression normalized to α-tubulin and shCont is indicated. **(B)** Analysis of tRNA expression in T47D parental and shMaf1-58 cells with R50 treatment. (**C**) Relative Bru incorporation by tRNA genes occupied by POLR3A/PR, POLR3A alone, or none of the three factors from R50 ChIP-seq data. (**D**) Relative Bru incorporation by tRNA transcripts of all high confidence tRNA genes from tRNAscan-SE in T47D or T47D shMaf1 cells.

## DISCUSSION

Studies on ER and PR gene regulation in breast cancer have overwhelmingly focused on genes transcribed by RNA Polymerase II (Pol II). While it is well established that the activity of various Pol III-transcribed genes increases in breast and other cancers, likely driven by oncogenes such as c-myc or activation of the PI3K-AKT-mTOR pathway (12,13), hormonal regulation of Pol III remains virtually unstudied. In breast cancer cells, PR is generally considered to be a modulator of ER transcription. Unliganded PR are predominantly nuclear and have been observed at active ER transcription complexes. In the presence of PR ligands, both receptors relocate to PR target genes (8,9), typically resulting in a more cytostatic or tumor suppressive phenotype. This effect is thought to involve reduced expression of ER-targeted mitogenic genes, although other mechanisms may also contribute. We previously observed that PR, but not ER, associates with Pol III at tRNA genes in breast tumors (10). In the current study, we describe a novel interaction between PR and the Pol III repressor Maf1 at tRNA genes in breast cancer cells, associated with decreased expression of target tRNAs. These findings suggest a broader role for PR in regulating cell growth and differentiation that operates independently of direct interference with ER-mediated transcription. This suggests the intriguing idea that PR-mediated regulation of select tRNA genes, and qualitative alterations in the tRNA population, may ultimately alter the translation efficiency of select mRNAs.

According to the genomic tRNA database (GtRNAdb), the human genome (hg38) contains 429 high-confidence tRNA genes (30). These encode 49 tRNA isoacceptors responsible for translating all 21 amino acids, including selenocysteine. Our analysis of T47D breast cancer cells revealed that the majority (88%) of tRNA loci were occupied by Pol III, as determined by POLR3A ChIP-seq. Interestingly, PR associated with approximately half of the Pol III-occupied tRNA genes in a progestin-dependent manner, spanning all 21 amino acids and all but one of the 49 codons. Consistent with our previous findings in UCD4 PDX tumors (10), these results suggest a robust pattern of PR occupancy across diverse tRNA codons. Changes in individual tRNA species or the tRNA pool can significantly impact mRNA translation elongation rates and the translation of specific mRNA transcripts. For example, overexpression of the initiator methionine tRNA (tRNA-iMet) in breast epithelial cells enhances proliferation and metabolic activity (33). Moreover, specific tRNAs such as tRNA-Arg-CCG and tRNA-Glu-UUC have been linked to increased metastatic potential in triple negative breast cancer cells, likely due to enhanced translational efficiency of pro-metastatic genes (34). Post initiation translational efficiency profoundly influences the proteome and plays a critical role in oncogenic properties (35–37). Codon usage, a major determinant of translation efficiency and mRNA stability, is intricately tied to cellular proliferation state and predicted to impact cancer progression, although the underlying mechanisms remain poorly understood. Certain KRAS-driven cancers, for example, arise from rare codon usage patterns (38). In our study, progestin treatment induced a widespread decrease in tRNA transcription which could have several implications. First, a general reduction in tRNA availability might contribute to slowed cell growth. Second, alterations in the tRNA pool could selectively promote or inhibit the translation of specific transcripts via codon usage biases. We speculate that progestin-induced changes in the tRNA pool may skew cellular processes towards differentiation rather than proliferation. Notably, the mechanism by which PR associates with tRNA genes remains unclear; motif analysis of PR-bound tRNA genes did not identify DNA sequences resembling conventional steroid receptor response elements. These results suggest that PR may engage these loci through an alternative, non-canonical binding mechanism.

In our studies, Maf1 exhibits a both cytoplasmic and nuclear localization with a fraction in the cytoplasm independent of progestin treatment (Supplementary Figure S5B) consistent with previous mammalian studies (41,42). In liver cancer cells, Maf1 levels are post-transcriptionally controlled via mTOR-mediated phosphorylation at Ser75 followed by ubiquitination; loss of Maf1 correlates with chemoresistance (43). Given progestins rapidly activate kinase signaling pathways (44), it is plausible that they impact Maf1 phosphorylation, although this was not assessed in our study. In yeast, the predominant mechanism of Maf1-mediated repression involves disrupting Brf1 and TFIIIB, thereby impeding Pol III recruitment (22,31,45). However, our study using short-term (1 h) progestin treatment observed no significant change in Brf1 occupancy between control and progestin-treated cells. Instead, Brf1 occupancy appears stabilized, as evidenced by increased ChIP peak amplitudes. Therefore, we propose a model whereby progestins recruit PR and Maf1 to specific Pol III-occupied tRNA genes, stabilizing the TFIIIB-Pol III complex. Orioli et al (32) previously proposed that Pol III might transiently occupy a gene in an unproductive state in human fibroblasts, although they did not investigate a potential role for Maf1. Our findings support a model in which progestins induce the formation of transcriptionally inactive PR-Maf1-POLR3A-Brf1 complexes at tRNA genes.

Interestingly, we find that Maf1 was also detected at select Pol II-transcribed genes with association augmented by progestin treatment (Supplementary Figure S6). This finding aligns with two studies that demonstrate occupancy of Maf1 in proximity to Pol II-transcribed genes in human cells. Thus, emerging data suggests broader roles for Maf1 beyond Pol III regulation in mammalian cells. In glioblastoma cell lines, Maf1 acts as a negative regulator of Pol I and Pol II transcription, potentially through its ability to regulate the TBP promoter (20,39). Moreover, Maf1 acts as a downstream effector of the tumor suppressor PTEN, inhibiting lipogenic gene expression and intracellular lipid accumulation in cancer cells (40). In this case, Maf1 was shown to repress Pol II transcription directly by its recruitment to the FAS promoter. Given our initial results, future studies may determine how Maf1 occupancy of RNA Pol II-dependent genes is regulated through PR.

This study has several limitations worth noting. First, to robustly investigate PR’s impact on Pol III activity, we selected breast cancer cells that are ubiquitously rich in steroid receptors. In contrast, normal mammary tissue cells are organized with punctate expression in the luminal epithelial cell layer. Exploring how PR impacts Pol III activity in this context could provide insight into how progestins drive mammary gland expansion and contraction through mammary stem cells (46,47). Second, while ChIP-seq assays offer a snapshot of PR chromatin binding, they do not fully capture the dynamic process of association and disassociation. Thus, PR may dynamically occupy enough tRNA genes at any given time to slow Pol III activity. This hypothesis gains support from the absence of traditional PREs near PR-occupied peaks, indicating direct recruitment to Pol III complexes. Additionally, our Maf1 ChIP results may not fully capture all binding sites, potentially underestimating its presence at some tRNA loci. Third, due to the stable tertiary structure and extensive modifications, there are no standardized high-throughput assays to quantitate individual pre-tRNA transcript levels. Various tRNA-sequencing methods have been reported (48–51), each with technical limitations and none yet adopted as standard practice in the field. Here we demonstrate that Bru-seq is a robust and relatively straightforward method for quantifying direct effects on pre-tRNA transcription. Finally, cancer cell genomes commonly exhibit irregularities, including chromosomal gains and losses that vary between tumors. Consequently, the number of tRNA genes likely differs among cancer types and individual tumors. In our investigation of two breast cancer models, we observed Pol III occupancy at tRNA genes encoding all 49 isoacceptors, confirming that the full set is collectively present in breast cancer cells.

In summary, our findings demonstrate that PR associates with and rapidly modulates Pol III transcription of tRNAs in breast cancer cells in a progestin-dependent manner. Progestins recruit the Pol III repressor Maf1 alongside PR at tRNA genes without disrupting Brf1 or POLR3A binding to DNA, suggesting a novel mechanism of Pol III repression by steroid receptors. These results have important implications for understanding steroid-mediated regulation of cell growth and differentiation across various tissues and cancers. This is particularly significant given most prior studies on Pol III have been conducted in systems lacking steroid hormone receptors. Notably, ChIP assays did not detect ER near tRNA genes in breast cancer cells. However, given that ER can regulate many genes via long-range chromatin looping (52), it remains plausible that ER could influence tRNA transcription through this mechanism, or indirectly through oncogenes such as myc. The identification of the tumor suppressor Maf1 as a potential co-repressor for PR and potentially other steroid receptors also highlights opportunities to modulate its activity through hormone signaling pathways.

## DATA AVAILABILITY

ChIP-seq data generated in this study have been deposited in the NCBI’s GEO database under the accession codes GSE283978 and GSE283979.

## SUPPLEMENTARY DATA

Supplementary Data are available at NAR Online.

## ACKNOWLEDGEMENTS

We thank the University of Colorado Cancer Center Genomic, Cell Technologies, and Biorepository Shared Resources supported by P30CA046934 for their technical assistance and services.

