## Supplementary figures for "Maf1 Cooperates with Progesterone Receptor to Repress RNA Polymerase III Transcription of Select tRNAs"

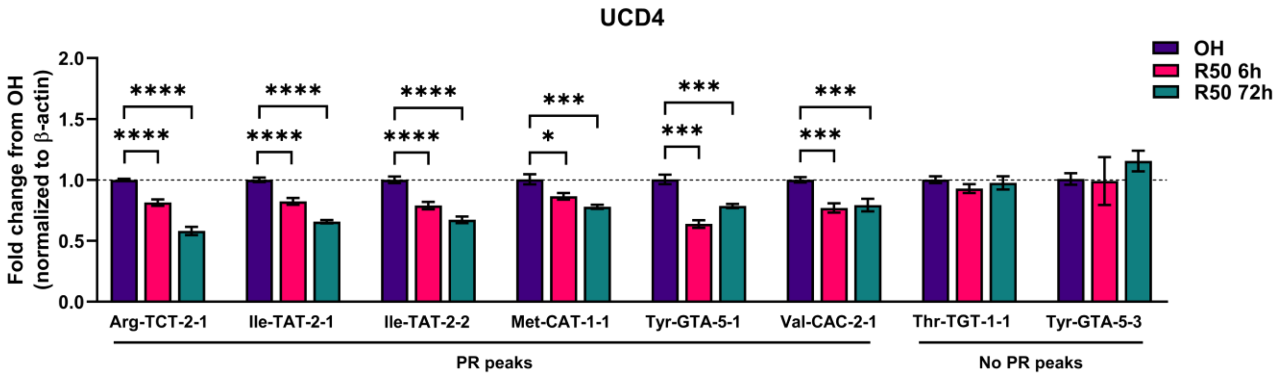


**Supplementary Figure S1. Progestin treatment downregulates pre-tRNAs.** qPCR analysis of pre-tRNA transcripts in UCD4 cells treated with vehicle (OH) or 10 nM R50 for 6 or 72 h. At least six biological replicates were used for analysis. Error bars depict mean +/-SEM. Significance was determined using Student's t test. * p<0.05, ** p<0.01, *** p<0.001, **** p<0.0001.

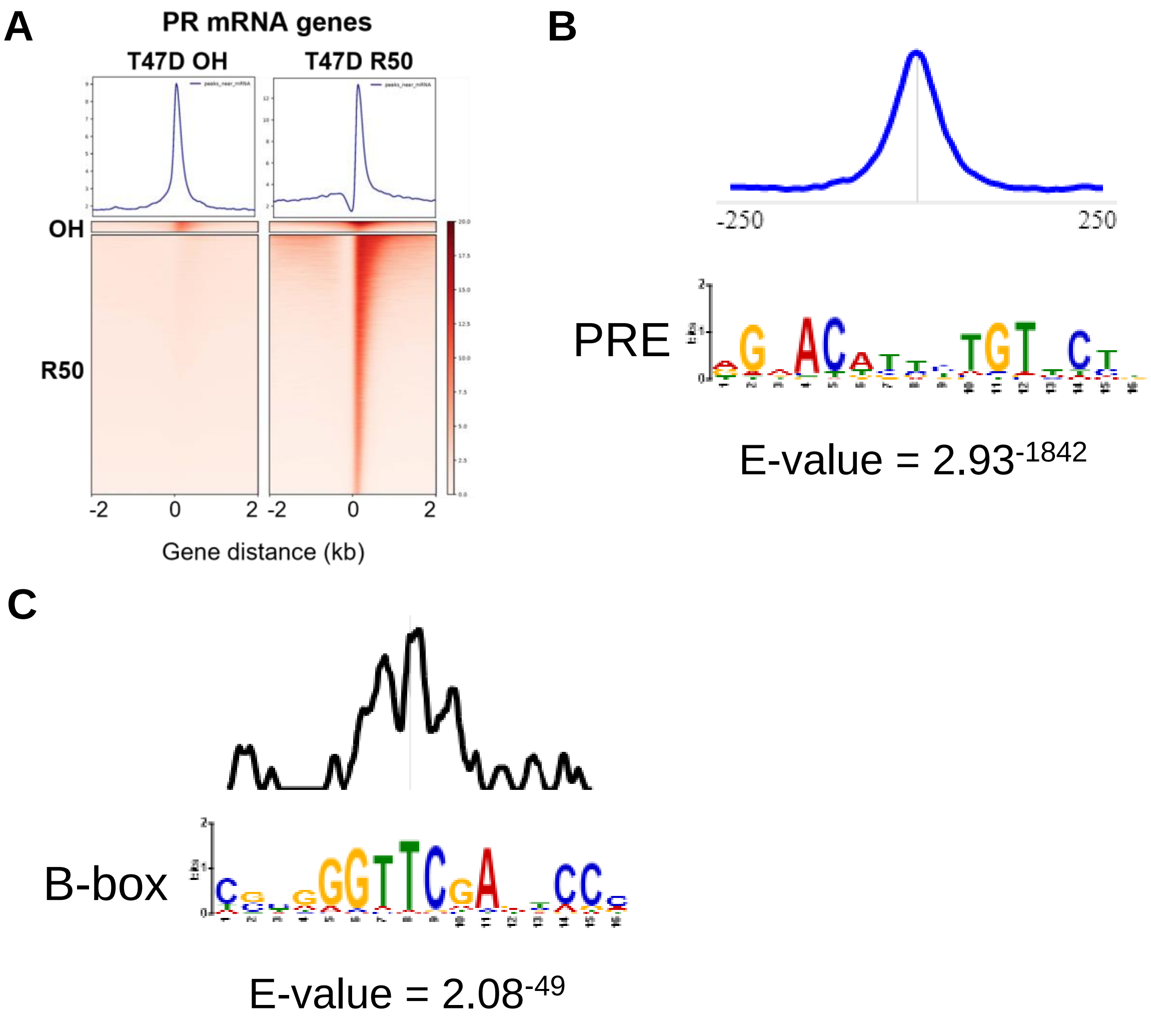


**Supplementary Figure S2. PR binding sites differ between mRNA and tRNA genes. (A)** Average signal plots and heatmaps for ChIP-seq binding events over mRNA genes in a horizontal window of +/- 2 kb with antibodies to PR in T47D cells treated with ethanol vehicle (OH) or 10 nM R50 for 1 h. Experiment was in biological triplicates. **(B)** Motif analysis of R50 PR all gene peak sequences identified sequences and sequence distribution consistent with PREs. **(C)** Motif analysis of R50 PR tRNA gene peak sequences identified the sequence and distribution consistent with a B-box. E-values were calculated using MEME-suite AME analysis.

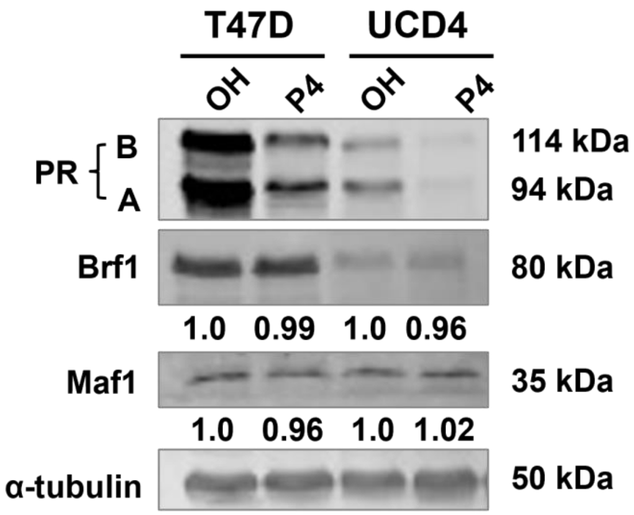


**Supplementary Figure S3. Maf1 and Brf1 protein levels do not change with R50 treatment.** Immunoblots for PR, Brf1, Maf1, and α-tubulin loading control in whole-cell lysates of T47D and UCD4 cells treated with vehicle (OH) or 10 nM R5020 (R50) for 24 h. UCD4 cells were pretreated with 10 nM estradiol for 48 h prior to OH/R50 treatment. Ratio of Maf1 or Brf1 expression normalized to α-tubulin is indicated.

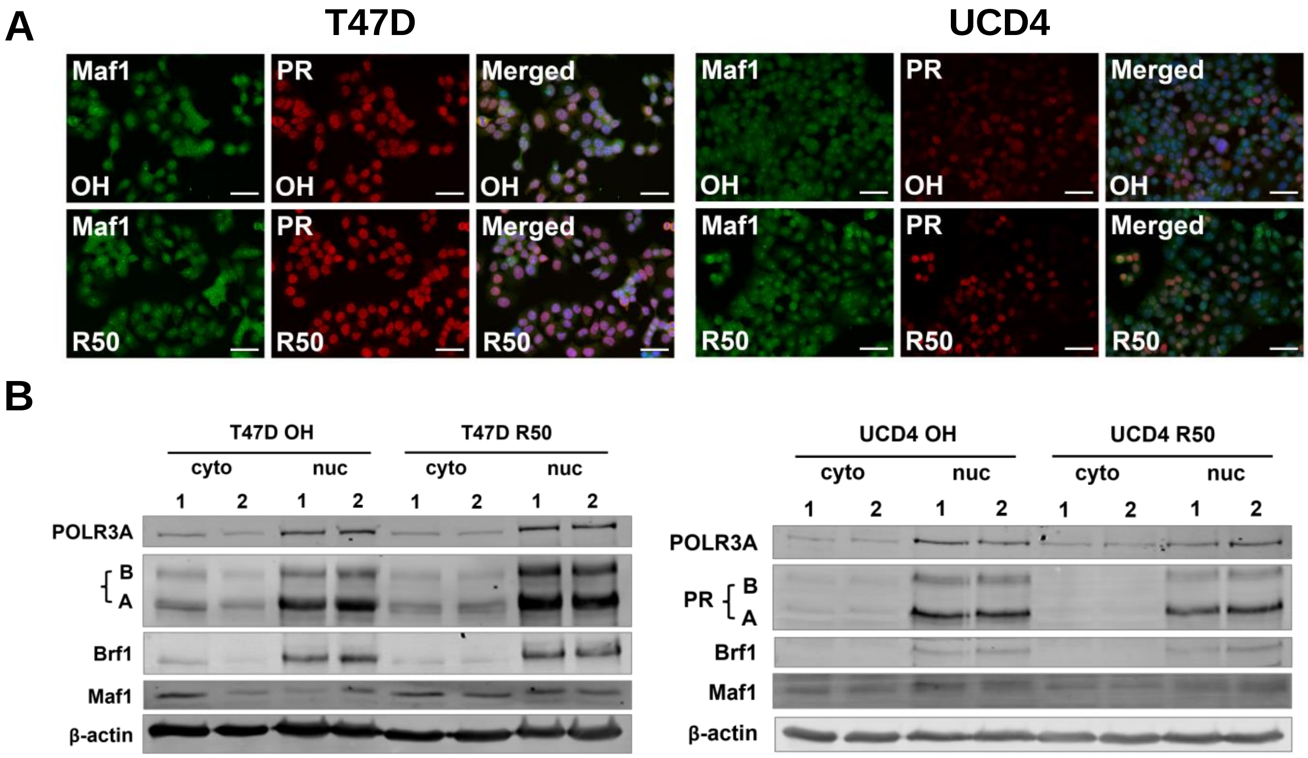


**Supplementary Figure S4. Maf1 co-localizes with PR. (A)** Immunocytochemistry for Maf1 and PR in T47D and UCD4 cells after 1 h of vehicle (OH) or 10 nM R5020 (R50) treatment. Maf1 (green), PR (red), and merged images plus DAPI counterstain (blue) are depicted. Scale bars, 50 microns. **(B)** Immunoblots for POLR3A, PR, Brf1, Maf1, and β-actin loading control in cytosolic and nuclear fractions of T47D and UCD4 cells treated with vehicle or 10 nM R5020 (R50) for 1 h.

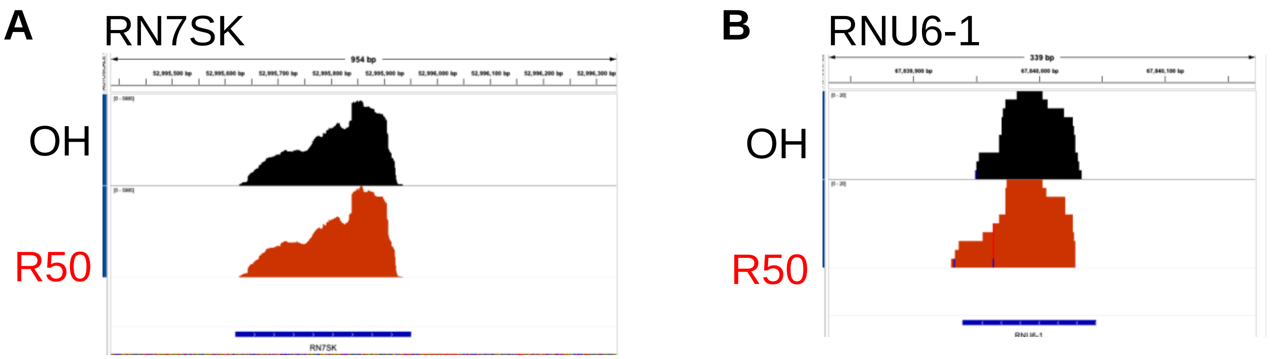


**Supplementary Figure S5. Nascent transcription from representative genes using Pol III type 3 promoters does not change with progestin treatment.** Bru-seq analysis in T47D cells treated with vehicle (OH) or 10 nM R5020 (R50) for 1 h. Relative Bru incorporation by **(A)** RN7SK and **(B)** RNU6-1 transcripts.

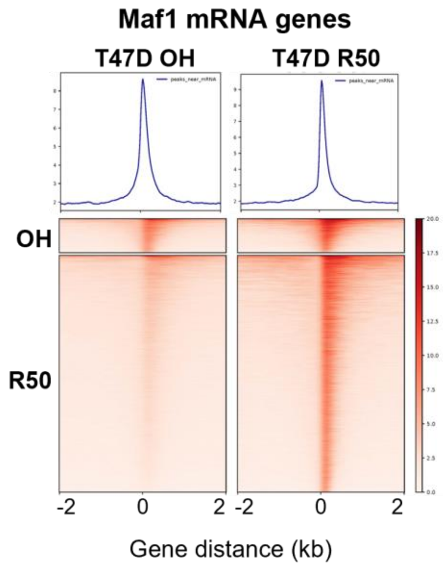


**Supplementary Figure S6. Maf1 occupies mRNA genes.** Average signal plots and heatmaps for ChIP-seq binding events over mRNA genes in a horizontal window of +/- 2 kb with antibodies to Maf1 in T47D cells treated with ethanol vehicle (OH) or 10 nM R50 for 1 h. Experiment was in biological triplicates.
